# Copiotrophs dominate rhizosphere microbiomes and growth rate potential is a major factor explaining the rhizosphere effect

**DOI:** 10.1101/2022.11.24.517860

**Authors:** José L. López, Nikolaos Pappas, Sanne WM Poppeliers, Juan J. Sanchez-Gil, Arista Fourie-Fouche, Ronnie de Jonge, Bas E. Dutilh

## Abstract

The structure and function of the root microbial community is shaped by plant root activity, enriching specific microbial taxa and functions from the surrounding soil as the plant root grows. Knowledge of bacterial rhizosphere competence traits are important for predictive microbiome modeling and the development of viable bioinoculants for sustainable agriculture solutions. In this work we compared growth rate potential, a complex trait that recently became predictable from bacterial genome sequences, to functional traits encoded by proteins. We analyzed 84 paired rhizosphere- and soil-derived 16S rRNA metabarcoding datasets from 18 different plants and soil types, performed differential abundance analyses and estimated growth rates for each bacterial genus. This analysis revealed that bacteria with a high growth rate potential consistently dominated the rhizosphere. Next, we analyzed the genome sequences of 3270 bacterial isolates and 6707 MAGs from 1121 plant- and soil-associated metagenomes, confirming this trend in different bacterial phyla. We next investigated which functional traits were enriched in the rhizosphere, expanding the catalog of rhizosphere-associated traits with hundreds of new functions. When we compared the importance of different functional categories to the predicted growth rate potential using a machine learning model, we found that growth rate potential was the main feature for differentiating rhizosphere and soil bacteria, revealing the broad importance of this factor for explaining the rhizosphere effect. Together, we contribute new understanding of the bacterial traits needed for rhizosphere competence. As this trait may be inferred from (meta-) genome data, our work has implications for understanding bacterial community assembly in the rhizosphere, where many uncultivated bacteria reside.

## Introduction

Soils represent the most complex and diverse microbiomes in the world. A notable extension of these is the rhizosphere, comprising the soil region near plant roots, which are influenced by root exudates and rhizodeposition [1]. The rhizosphere microbiome has been studied with the aim of harnessing plant-microbe interactions and improving sustainable agriculture [1]. Rhizosphere microbiomes assemble by recruiting a subset of the microbiota present in the soil surrounding plant roots, also known as the bulk soil, and these recruited microbes may pose beneficial, neutral, or detrimental effects on the host plants [1, 2]. The changes in microbial community composition that take place from the soil towards the root have been dubbed the rhizosphere effect [3]. It involves a decrease in species richness imposed by stronger selection on microbes towards the roots into sequential regions known as the rhizosphere, rhizoplane, and endosphere [4]. The strength of the rhizosphere effect varies extensively between different studies [4,5-9,10]. While methodological differences between these studies, including rhizosphere and microbiome isolation cannot be ruled out, there may also be physiological factors that influence the extent of this effect [3]. Among other factors, the rhizosphere effect may be influenced by the host plant species [11], the stage of the plant life cycle [12], or the location on the root [13].

Understanding the biological signatures that allow a microbe to colonize and thrive in the rhizosphere has been a complex and intricate endeavor, and it remains an open question which key factors drive rhizosphere community assembly [3]. Approaches to understand rhizosphere competence have frequently included analyzing compositional or functional changes between root-associated bacteria and soils, revealing that some functions or taxa are enriched in different rhizosphere microbiomes [14-18]. Others have identified core microbes that are consistently present in the rhizosphere across different plant hosts and conditions [2, 19]. Still, the variation in, and diversity of rhizosphere microbiomes described in different studies makes it difficult to make general statements on the bacterial traits that are common determinants of rhizosphere competence in different plants.

Besides descriptions of the changes in taxonomic or functional composition of the rhizosphere microbiome, alternative genomic signatures for the rhizosphere effect are not frequently explored. One relevant ecological aspect to describe microbial lifestyle is growth rate potential. There have been indications that shifts in the soil microbial composition in response to carbon availability can to some extent be predicted in terms of copiotrophs, microorganisms adapted to high nutrient conditions with faster growth rates, and oligotrophs, microorganisms adapted to low nutrient conditions with slow but more efficient growth [20]. Similarly, copiotrophs are enriched in soils with higher presence of labile organic substrates (i.e., glycine, sucrose), while oligotrophs are enriched in soils containing recalcitrant chemicals (i.e., cellulose, lignin, or tannin–protein) [21]. Finally, an increase in the relative abundance of copiotrophic bacteria in soils has been associated with elevated nitrogen and phosphorus agricultural inputs [22]. Together, these observations indicate that compounds secreted by plant roots could contribute to the selection of copiotrophs in the rhizosphere, thus playing a role in establishing the rhizosphere effect.

Recently, estimations of maximal growth rate, further referred to as growth rate potential, have become possible from genomic data, without the need of culturing [23]. These predictions are based on a model that computes signals including the codon usage bias in a genome, codon usage pattern consistency in highly expressed genes, and genome-wide codon pair bias. The model provides an estimation of a minimum doubling time of an organism and allows for the classification of a bacteria into copiotroph or oligotroph [23]. Being applicable to bacteria, archaea, as well as eukaryotic microbes, growth rate potential has been explored recently using this model-based approach in different biomes, such as in oligotrophic marine systems or the nutrient-rich human gut [24]. In a recent work this approach was also used in whole communities of marine samples [25], where the authors found a decreasing community-wide average growth rate potential correlated with depth, probably owed to a decrease in nutrient availability. This suggests that nutrient gradients may affect the growth rates in different biomes, the rhizosphere being one natural habitat where to test this hypothesis. Here, we investigated whether microbial growth rate potential is a predictor of rhizosphere enrichment. We analyzed 460 rhizosphere and 232 bulk soil 16S rDNA metabarcoding samples comprising 84 paired rhizosphere and bulk-soil datasets from 18 different plant genotypes, a set of previously analyzed isolated genomes from plants and soils [14], and MAGs (metagenome-assembled genomes) from 501 rhizosphere and 620 soil full metagenomes from diverse studies to analyze how growth rate potential may contribute to the rhizosphere effect.

## Methods

All R scripts and Jupyter Notebooks for analysis, plotting and tables used in this analysis are available on GitHub (https://github.com/JoseLopezArcondo/rhizosphere_microbial_growthRates). All visualizations were done using ggplot2 [26], ggpubr [27], and edited using InkSkape [28].

### Matching rhizosphere and bulk soil datasets in 16S rDNA metabarcoding datasets

We selected metabarcoding projects from the MGnify database [29] based on i) having both rhizosphere and associated bulk soil samples available, and ii) having sufficient sequencing depth, i.e., with more than 10000 reads per sample (Supplementary Table 1). Biom files corresponding to SSU rRNA OTU counts and their taxonomic assignments were downloaded. We prepared abundance matrices at the genus rank containing per sample the sum of amplicon sequence variant (ASV) counts of all ASVs per given genus, and DESeq2 [30] analysis was performed to identify genera enriched in the rhizosphere or in bulk soil (adjusted *p*-values < 0.05, Benjamini-Hochberg FDR method).

### Minimal doubling time predictions

To estimate growth rate potential of bacterial genera, we used the estimated growth rates from the gRodon online (EGGO) database. Based on the genera identified above, we first collected the predicted minimum doubling time (PMDT) of all genomes belonging to these genera from the EGGO database [23], and then calculated the median PMDT (mPMDT) per genus. The PMDT of metagenome-assembled genomes (MAGs, see details below) and isolated genomes was estimated using the gRodon R package version 1.0.0 [23]. To this end, ribosomal protein-coding genes were obtained by searching “rps”, “rpm”, “rpl” terms in the “Preferred_names’’ column of the eggNog annotation file (see details below). Ribosomal protein-coding genes were assumed to be highly expressed [31] and therefore used as a reference gene set for gRodon analysis. As parameters, we used ‘partial’ mode, and ‘vs’ training set. Genera, isolated genomes, and MAGs were considered as copiotrophs or oligotrophs using a cutoff of PMDT < 5hs and PMDT ≥ 5hs, respectively, according to Weissman et al. [23].

### Functional and phylogenetic annotation of metagenome-assembled genomes (MAGs)

We retrieved MAGs of medium and high quality (MQ-HQ) from rhizosphere and soil metagenomes from the Integrated Microbial Genomes and Microbiomes (IMG/M) database [32]. These MAGs were generated by the IMG/M pipeline [32], using Metabat v2:2.15 [33] and checkM v1.1.3 [34], and were classified as medium quality (MQ) or high quality (HQ) according to the Genomic Standards Consortium criteria [35], i.e., genomes with completeness estimates of ≥ 50% and less than 10% contamination. To obtain the contigs that constituted each MAG, we downloaded the assembled metagenomic contigs along with extensive functional annotations. Using mapping files from IMG/M connecting the contig IDs to MAGs we isolated sequences and annotations for each individual MAG. We then used individual MAG annotations consisting of Clusters of Orthologous Groups (COGs), KEGG Orthology (KO) and Protein Families (Pfam) to create binary matrices, consisting of the presence or absence of a specific function in each MAG. For gRodon predictions on MAGs and isolated genomes, nucleotide and protein sequences for genes were predicted using Prodigal version 2.6.3 [36] and functions were annotated using eggNog mapper version 2.1.3 [37].

To interpret the evolutionary relationships of the recovered MAGs in the context of known strains, we generated a maximum-likelihood phylogenetic tree based on concatenated GTDB marker genes, using GTDB-Tk version 1.3.0 and GTDB database release 95 ([38], gtdbtk identify, align and infer commands).

### Phylogeny-aware functional enrichment analyses

To identify functions associated with rhizosphere or soil bacteria, MAGs were labeled as belonging to rhizosphere or soil, and we constructed a vector of binary target labels, corresponding to rhizosphere (1) and soil (0), based on the type of metagenomic sample where it was recovered from (see Supplementary Table 2). Functional binary matrices including the presence or absence of each functional category were used as independent variables and, together with the bacterial phylogenetic tree, were used as input in phylogeny-aware functional enrichment analyses using Phylogenetic Generalized Linear Models (PhyloGLMs) [39] from the phylolm R package v. 2.6.2. Similar models were generated to identify enriched functional groups in MAGs labeled as copiotroph or oligotroph, based on their PMDT. For PhyloGLM analyzes we labeled copiotrophs (PMDT < 5hs) and oligotrophs (PMDT ≥ 5hs), see Supplementary Table 3. We corrected p-values (p-adjust) with Benjamini-Hochberg FDR and used p-adjust values < 0.05 to consider significantly enriched functions.

### Machine learning models

Based on COG, Pfam and KO functional matrices for MAGs, we constructed binary feature matrices (presence/absence of each functional ortholog) with the additional feature copiotroph (1) - oligotroph (0) based on PMDT < 5hs and PMDT ≥ 5hs, respectively. Also, we constructed a vector of binary target labels, corresponding to rhizosphere (1) and soil (0), depending on whether most of the MAGs from the genus were obtained from rhizosphere or bulk soil metagenomes, respectively. We formulated Random Forest (RF) models and Gradient Boosting Classifier (GBC) models to classify whether a MAG was associated with rhizosphere or soil. The final datasets consisted of 6707 MAGs (3692 from rhizosphere and 3015 from soils) with 8680, 4841, and 9132 features for KO, COG, and Pfam-based binary matrices, respectively. With the goal of identifying which functional features were most important for rhizosphere competence, we trained and evaluated machine learning models with the scikit-learn Python package (https://scikit-learn.org/). We used 5-fold cross validation to verify the models, and tested different parameter settings for RF models, including number of trees in the forests (n_estimators), maximum number of features in each node (max_features), maximum depth of trees (max_depth), maximum leaf nodes in the trees (max_leaf_nodes), minimum number of samples to create a leaf node (min_samples_leaf), minimum number of samples to generate a split (min_samples_split), and for GBC models we evaluated different settings of n_estimators, max_depth, and the learning rate of subsequent trees (learning_rate) parameters. Finally, Gini feature importances, which are a measure of the relative accumulation of the impurity decrease for each feature in the model, were obtained, and the main features were analyzed with STRING version 11.5 [40].

## Results

### Rhizosphere bacteria have shorter predicted doubling times than soil bacteria

To investigate whether growth rate potential predictions correlate with rhizosphere enrichment, we re-analyzed previously published metacommunities, comprising plant rhizospheres and their associated bulk soils. These include microbiomes associated to 18 different plant genotypes and conditions such as: *Arabidopsis thaliana* ecotypes and sister species [9], *A. thaliana* Col-0 ecotype under light-dark cycles [41], wild and modern accessions of *Phaseolus vulgaris* (common bean) [42], *Zea mays* grown in soils with different crop rotation systems [43], *Sorghum bicolor* under drought stress and control conditions, at different timepoints in their lifecycle [44], and *A. thaliana* Col-0 sampled at different stages along a bulk soil-to-rhizosphere gradient [45]. First, we obtained metacommunity data at the amplicon sequence variant (ASV)-level from diverse sources (see Materials and Methods), and then grouped ASVs at the genus level. Next, we performed differential abundance analysis, identifying which genera were significantly enriched in rhizospheres and bulk soils (p-adjust < 0.05, log-2 fold change (L2FC) > 1 for soil-enriched genera, L2FC < -1 for rhizosphere-enriched genera). We then mapped each genus to the EGGO database containing the predicted growth rates from hundreds of thousands publicly available genome sequences, and assessed the growth rate potential by using the median predicted minimal doubling time (mPMDT) of genomes belonging to these genera. This allowed us to compare the mPMDT of rhizosphere-enriched to soil-enriched genera (see Materials and Methods), which revealed that rhizosphere-enriched genera have on average faster growth rates (lower mPMDT) than soil-enriched genera (Figure 1 a-b), consistently across different experimental conditions.

**Figure 1.**
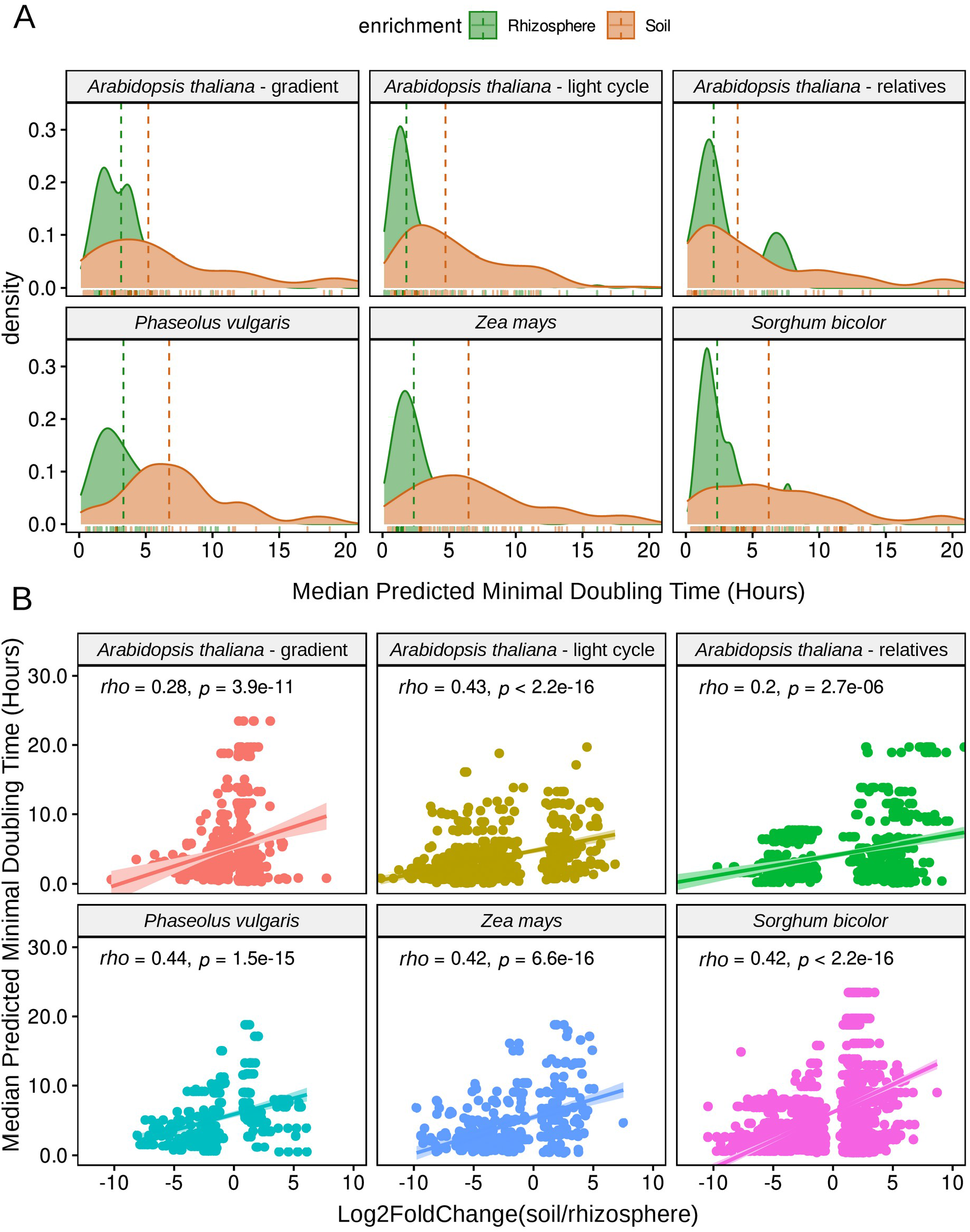
Median predicted minimum doubling times (mPMDT) of bacteria enriched in rhizospheres are lower than those in associated bulk soils. DESeq2 log2 fold-change was used to categorize bacteria as being enriched in the rhizosphere (L2FC < -1) or soil (L2FC > 1). Soil-enriched bacteria tend to have a higher mPMDT. **A.** Density distribution of bacteria enriched in the rhizosphere or soil. **B.** A positive correlation exists between soil enrichment and mPMDT, i.e. the rhizosphere contains faster growers than the bulk soil.

First this general trend was observed in different species of the same plant genus (*Arabidopsis*), in modern and wild accessions of a same species (*P. vulgaris*), and in different plant hosts (*A. thaliana, P. vulgaris, S. bicolor* and *Z. mays*) (Supplementary Figures 1-3), showing that although plant hosts induce specific compositional shifts in the rhizosphere microbiomes [11], faster growth rates to colonize rhizosphere seems to be a common factor. Second, although the host’s circadian rhythm induces changes in the rhizosphere microbiome and the soil organic matter composition [41], it does not affect the rhizosphere-enrichment of copiotrophs (Supplementary Figure 2). Third, different soil conditions, such as crop rotations and drought stress do not modify this general trend either, as shown here in *Z. mays* and *S. bicolor* ([43, 44], Supplementary Figures 1,3). Finally, in a study where a gradient from bulk soil to the rhizoplane was experimentally dissected and analyzed separately [45], we observed that copiotrophs increased gradually as samples were taken closer to the root (Supplementary Figure 4). Thus, we observed that the trend for fast-growing bacteria to colonize the rhizosphere was consistent and independent of the plant species or ecotype, soil type or experimental condition.

When we observed exceptions to this trend, i.e., copiotroph bacteria that were enriched in soils, these were mostly among Firmicutes (Supplementary Figures 1-3). Firmicutes are copiotrophs and are among the fastest growers in the bacterial tree [23], thus it may be harder to prove this trend in the narrower mPMDT distributions present in this particular phylum.

When analyzing the correlations between the L2FC enrichment scores and the mPMDT by the four most abundant phyla in our 16s rDNA data (Actinobacteria, Bacteroidetes, Firmicutes and Proteobacteria), we found a significant correlation between mPMDT and rhizosphere enrichment in *Proteobacteria* in all projects (Supplementary Table 1). When merging all samples from different projects, enrichment of the copiotrophs was significant for Proteobacteria, Actinobacteria, Bacteroidetes, Acidobacteria and Verrucomicrobia, but not in Firmicutes (Supplementary Figure 5). Thus, 16s rDNA data shows that copiotroph genera are preferentially enriched in rhizospheres compared to bulk soils in members of these main bacterial phyla.

### Bacterial genomes confirm copiotrophs are predominant in the rhizosphere

To further analyze changes in growth rate potential distributions using genome sequences of isolated bacteria, we estimated PMDTs in genomes from cultured bacteria isolated from plant, non-plant, root/rhizosphere and soil biomes reported in Levy et al. [14], including 3,270 genomes classified into taxonomic groups Actinobacteria groups 1 and 2 as defined by the authors, Alphaproteobacteria, Bacillales, Bacteroidetes, Burkholderiales, Pseudomonas and Xanthomonadaceae [14], by means of gRodon, which analyzes codon usage patterns in genes of each bacterial genome [23]. In Alphaproteobacteria and Bacteroidetes groups, bacteria isolated from rhizoplane and endophytic compartments (root associated, RA) have lower PMDT than those isolated from soils (Supplementary Figure 6). A similar observation extended to Actinobacteria_2, Alphaproteobacteria, Bacillales and Bacteroidetes, when comparing bacterial genomes isolated from plant niches, including rhizospheres (plant associated, PA) and from non-plant environments (NPA, which includes both soils and other environments like marine or clinical). Thus, although isolation protocols select for copiotrophs [23], we still observed an enrichment in copiotrophs in cultured isolates from plant environments when compared to those obtained from soil.

To avoid any possible biases associated with bacterial cultivation, we extended our genomic analyses to metagenome-assembled genomes (MAGs). We downloaded 501 whole-metagenome rhizospheric samples from different plants and 620 whole-metagenome soil samples from different biomes. We then recovered 3679 high-quality and medium-quality (HQ/MQ) bacterial MAGs from rhizospheres and 2784 HQ/MQ bacterial MAGs from soils (Materials and Methods, Supplementary Table 2). MAGs included members of the phyla Actinobacteriota (1453), Proteobacteria (including 1063 MAGs from Gammaproteobacteria, and 869 MAGs from Alphaproteobacteria), Acidobacteriota (850), Bacteroidetes (440), Patescibacteria (322), Verrucomicrobiota (248), Gemmatimonadota (200), Myxococcota (187), Planctomycetota (133), and Chloroflexota (127), Nitrospirota (76), Eisenbacteria (56), Methylomirabilota (51), Desulfobacterota (47), Desulfobacterota_B (45), and Firmicutes (36), among others (Supplementary Table 2). PMDT could be predicted for 6355 of these MAGs, of which 2629 were copiotrophs, 3726 were oligotrophs, while 3652 were obtained from rhizosphere samples, and 2703 were obtained from soils. A chi-squared test shows that PMDT and the isolation niche are significantly associated (Pearson’s Chi-squared, p-value < 2.2e-16, Supplementary Table 3). For a MAG, being copiotroph is significantly associated with colonizing the rhizosphere (PhyloGLM, estimate: 1.47, p-value: 2e-16; Figure 2).

**Figure 2.**
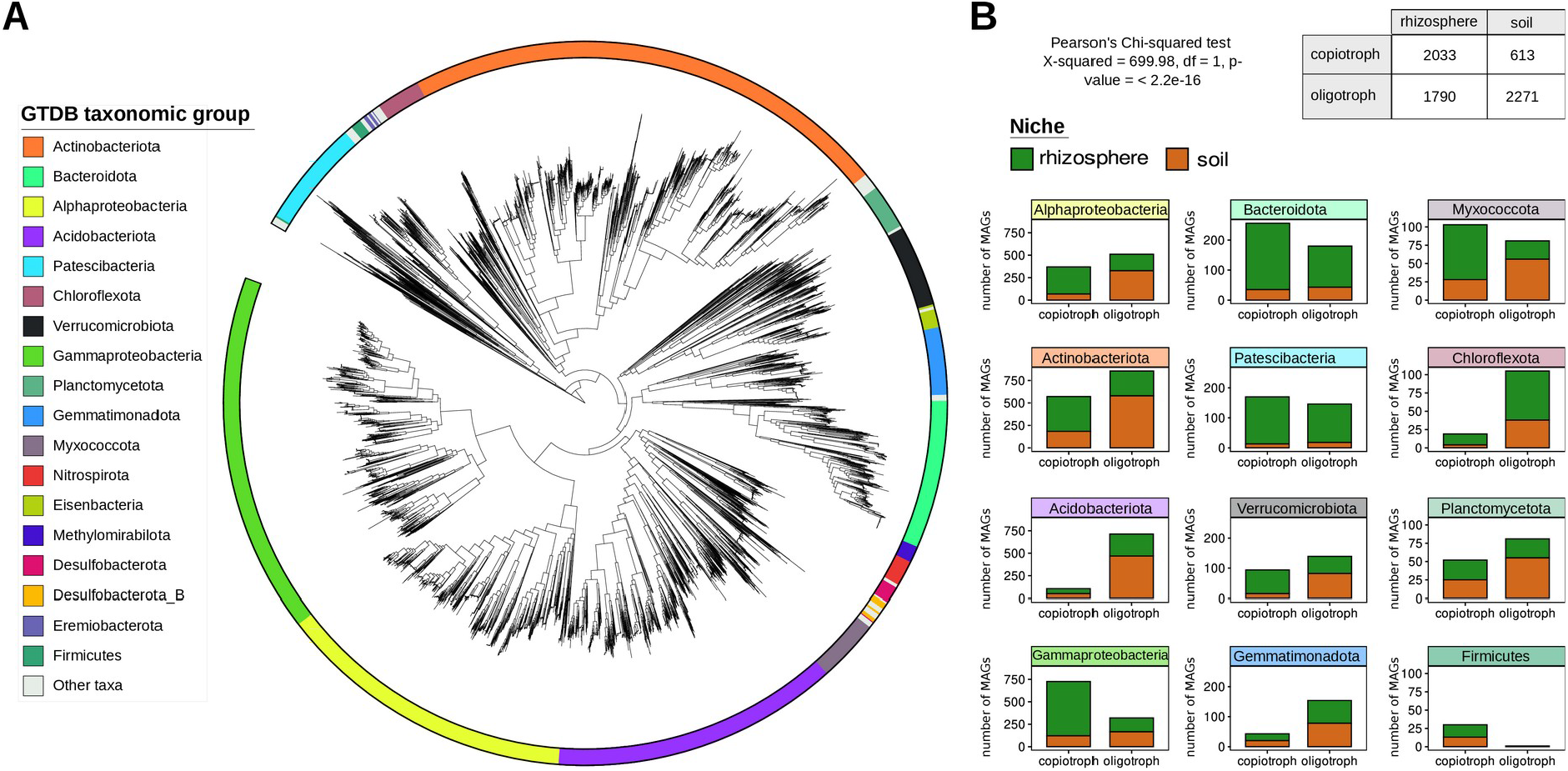
MAGs taxonomy, niche, and growth rate status. **A.** Unrooted maximum-likelihood phylogenetic tree inferred from multiple sequence alignments of GTDB bacterial marker genes from MAGs. The tree was generated with GTDB-Tk and displayed using iTol [46]. **B.** MAGs are classified according to their isolation biome and growth rate status (copiotroph or oligotroph) for each of the main GTDB taxa.

The analysis of MAGs allowed us to compare predicted growth rates of unculturable bacteria across a wide range of taxonomic groups (Supplementary Table 3). As shown in Weissman et al. [23], collections of isolates fail to capture the most slowly growing members of the communities, when compared to MAGs or single-amplified genomes (SAGs) from the same environments. Despite being obtained from diverse metagenomes and belonging to different plants and soils, our predictions of PMDT, as shown in Figure 3, revealed that MAGs obtained from rhizosphere metagenomes have significantly lower PMDT than MAGs obtained from soils across all major taxonomic groups except Firmicutes, which have an extremely low and narrow PMDT distribution.

**Figure 3:**
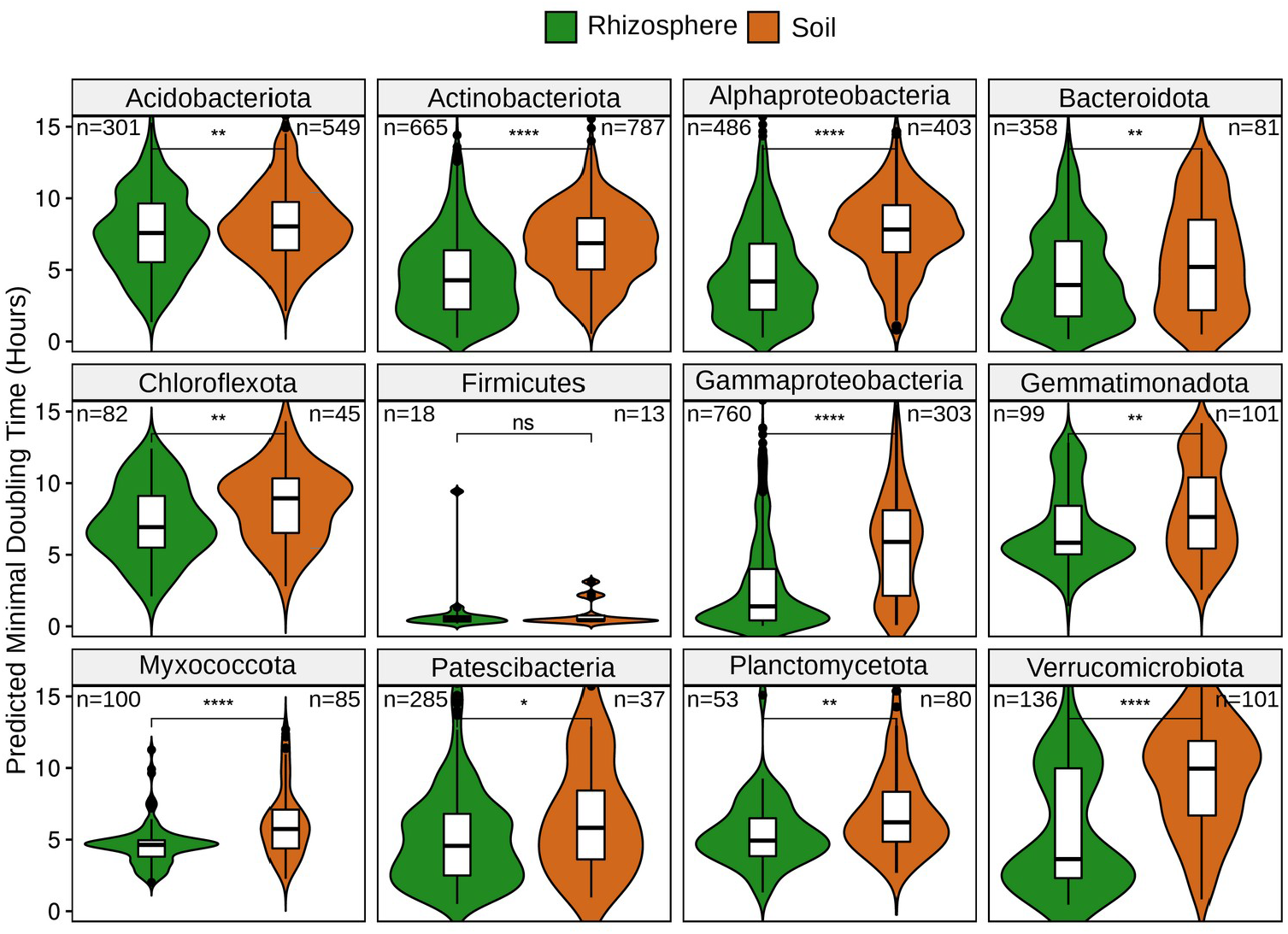
PMDT in MAGs from rhizosphere and soil metagenomes. Distributions of predicted minimal doubling times in MAGs from rhizosphere and soils were compared with Mann-Whitney test (ns: p > 0.05, *: p <= 0.05, **: p <= 0.01, ***: p <= 0.001, ****: p <= 0.0001).

### MAGs provide a catalog of functions associated with rhizosphere colonization and copiotroph / oligotroph lifestyles

To analyze which functions are significantly enriched when comparing MAGs from rhizospheres or soils and with copiotrophic or oligotrophic lifestyles, we employed a phylogenetic-aware approach (PhyloGLM) to compare genome functional content (KEGG orthology, KO). Figure 4A reveals that in Actinobacteria, Alphaproteobacteria, Bacteroidota, and Gammaproteobacteria, most functional categories were enriched in copiotrophs, while in Acidobacteria, most functional categories were enriched in oligotrophs. This highlights the differences in functional categories present in the genomes of copiotrophs and oligotrophs in these taxa. We then compared the genome size between the groups (estimated as gene counts per genome, Figure 4B) and found significantly larger genomes in copiotrophs from Actinobacteria, Bacteroidetes, and Gammaproteobacteria, and in oligotrophs in Acidobacteria, while no difference in genome sizes was found in Alphaproteobacteria, consistent with the enrichment of different functions in copiotrophs and in oligotrophs (Figure 4A). Interestingly, despite this difference in genome content, oligotrophs showed consistent enrichment in metabolism of terpenoids and polyketides, and metabolism of other amino acids, which include functions that are potentially relevant to the oligotrophic lifestyle. A similar pattern of genome content variation can be observed when comparing enriched processes in rhizosphere or soils in Acidobacteria and Gammaproteobacteria, although no significant differences in genome content were found in Bacteroidetes and Actinobacteria, and smaller MAGs were found in Alphaproteobacteria in rhizospheres, compared to those from soils. These patterns were also consistent with the enrichment of the different metabolisms in MAGs from rhizospheres or soils. Investigating why these differences in genome size exist in each taxonomic group and which functions are frequently missing in the smaller genomes could improve our understanding of copiotrophic and oligotrophic lifestyles.

**Figure 4.**
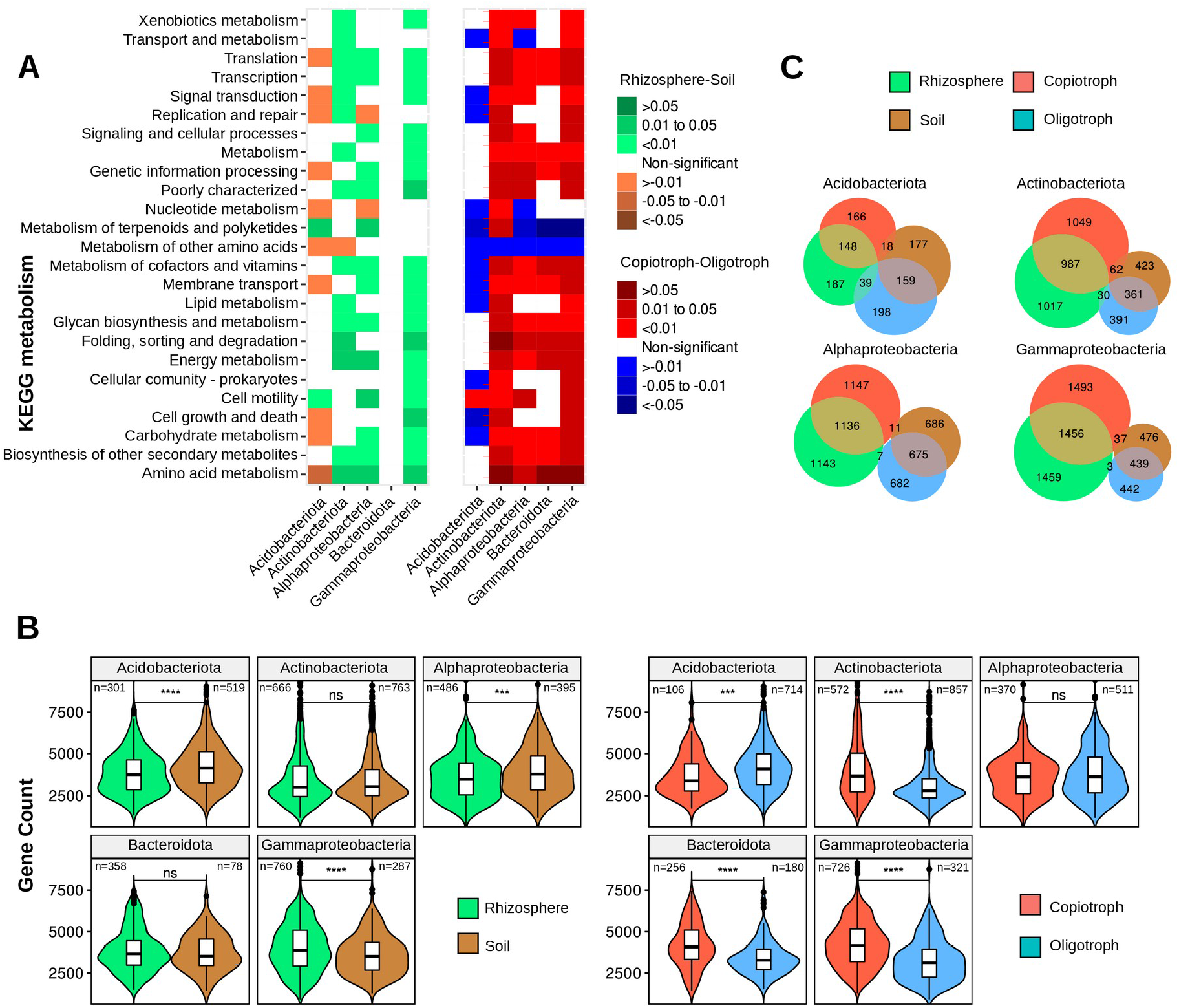
**A. Enrichment of KEGG functional categories in MAGs from 5 most representative taxa.** Differences in KO categories between rhizosphere and soil MAGs (left), and between copiotroph and oligotroph MAGs (right). Heatmaps indicate the level of enrichment based on the PhyloGLM test (p-adjusted values <0.05, Benjamini-Hochberg FDR method). **B. Gene Counts in MAGs from rhizosphere or soil (left) and copiotroph or oligotroph (right)**. Number of MAGs in each category is indicated, distributions of predicted minimal doubling times in MAGs from rhizosphere and soils were compared with Mann-Whitney test (ns: p > 0.05, *: p <= 0.05, **: p <= 0.01, ***: p <= 0.001, ****: p <= 0.0001). **C. Euler plots with significantly enriched KO functions in MAGs.** Plots show the number of enriched functions in the rhizosphere (green) or soil (brown), and copiotroph (red) or oligotroph (blue). Many more enriched functions were shared between rhizosphere-copiotroph and soil-oligotroph than between rhizosphere-oligotroph and soil-copiotroph.

When testing the enrichment of individual KO, COG and Pfam functions, we observed that many enriched functions overlap between rhizosphere-enriched bacteria and copiotrophs, and between soil-enriched bacteria and oligotrophs (Figure 4C, Supplementary Figures 7-10, Supplementary Table 4). A higher number of significantly enriched functions were obtained in most represented taxa (Supplementary Table 4), especially in Alphaproteobacteria, Gammaproteobacteria, Acidobacteria, and Actinobacteria.

### Growth rate potential is the main predictor of rhizosphere enrichment

To assess which functional features were important for copiotrophs or oligotrophs, we trained Random Forest (RF) and Gradient Boosting Classifier (GBC) models to predict the rhizosphere-or soil-association of a MAG based on a binary matrix including the presence or absence of KOs, COGs and Pfams, as well as its status as a copiotroph or oligotroph. We used Grid Search with Stratified Cross-Validation to evaluate how changing different parameters affected the RF and GBC models. We observed that increasing the number of trees (n_estimators) above 60 did not significantly increase the F1-score with neither COG, nor KO, nor Pfam matrices (Supplementary Figure 11). Also, changing the maximum number of features assessed at each node (max_features), or other pre-pruning parameters (see methods) did not result in a significant improvement of the RF models (Supplementary Figure 11). Thus, for the final model we set the number of trees to 300 and used default values for the remaining parameters, obtaining overall 5-fold cross-validated accuracy scores of 92.3%, 91.6%, and 91.7%, precision scores of 92.1%, 91.2%, and 91.4%, recall scores of 94.1%, 93.8%, and 93.8%, and F1-scores of 93.1%, 92.5%, and 92.6% for KO, COG, and Pfam based models, respectively. Moreover, we also trained GBC models and tuned parameters varying the number of trees (n_estimators, Supplementary Figure 12) with a grid combining different learning rates and maximum depth of trees (Supplementary Figure 13). We found optimal parameters at learning_rate=0.27, max_depth=14, and n_estimators=100 (Supplementary Table 5). With these parameters we obtained overall accuracy of 93.1%, 93.2%, and 92.8%, precision of 93.0%, 93.2%, and 92.6%, recall of 94.6%, 94.5%, and 94.4%, and F1-score of 93.8%, 93.8%, and 93.5% for KO, COG and Pfam based models, respectively.

With these classifiers in hand, we analyzed which features were important for classification of MAGs into the rhizosphere or soil categories. All 6 models (i.e. RF and GBC models based on KO, COG and Pfam binary matrices) identified the oligotroph/copiotroph status of a MAG as the most important feature, suggesting that a high growth rate potential is important for successful colonization of the rhizosphere (Supplementary Figure 14-15). Growth rate is a complex microbial trait, estimated here from codon usage patterns. It may be associated with some of the functional features that are also used as predictors in the models. However, our results clearly show that this trait is more important than any other individual function, highlighting the high predictive potential of this complex microbial trait that is readily inferred from the genome sequence.

### Rhizosphere and soil-associated functional traits

Besides growth rate potential, the machine learning models trained above also allowed us to rank the functional features by their importance to classify rhizosphere or soil bacteria. To further investigate this, we used the COG-based RF most important functions and filtered the ones that were also significant in at least one taxon in either soil or rhizosphere in the PhyloGLM analysis (Supplementary Table 6). We then searched for functional connections between the most important COGs in the STRING database [40]. Figure 5 shows that many of the COGs that were important for predicting rhizosphere-or soil-association have functional connections. One of the main clusters of rhizosphere-associated COGs contains functions involved with flagella. Flagellar motility has been reported as an important trait for rhizosphere colonization in other studies [47,48]. We observed that in PhyloGLM results, flagellar proteins were associated to rhizosphere and copiotrophs in Alphaproteobacteria, Acidobacteria, and Gammaproteobacteria, to copiotrophs in Actinobacteria, and never associated to soil or oligotroph, confirming the importance of active motility in rhizosphere colonization in the different taxa (Supplementary Table 4). Linked to this cluster we found an inter-membrane structural component of the type VI protein secretion system (T6SS, COG3521), which has been associated with modulating the microbial interactions and promoting rhizosphere competence of plant-beneficial bacteria [49, 50]. When we searched for the other COGs which compose the T6SS, we found that, when significantly enriched, in all cases they were associated either with rhizosphere or with copiotrophs, especially in Gammaproteobacteria (Supplementary Table 4). Second, we found a secreted acid phosphatase (COG2503), which may be involved in an adaptive response to low-phosphate stress in the rhizosphere [51]. Another connected cluster of COGs consists of proteins related to sugar catabolism, such as beta-galactosidase (COG3250, COG2731), alpha-L-fucosidase (COG3669), alpha-L-arabinofuranosidase (COG3534), beta-xylosidase (COG3507), a Na+/melibiose symporter or related transporter (COG2211), a DNA-binding transcriptional regulator of sugar metabolism of DeoR/GlpR family (COG1349), a mannose or cellobiose epimerase (COG2942), and a fructose/tagatose bisphosphate aldolase (COG0191), whose functions are implicated in mucilage polysaccharide degradation [52]. In a recent study in which the adaptation to plant colonization was tested in *Bacillus thuringiensis*, the authors found that metabolic pathways related to plant polysaccharides were upregulated in the adapted strain, including metabolism of various carbohydrates, such as cellobiose, pyruvate, and galactose [53]. Carbohydrate transport and metabolism functions are also overrepresented in copiotrophs [23]. Notably, the top five most important features in the RF model include a predicted sulfurtransferase (COG1054), an uncharacterized Zn-ribbon-containing protein (COG2824), an uncharacterized FAD-dependent dehydrogenase (COG2509), a tRNA A37 threonylcarbamoyladenosine dehydratase (COG1179), and an uncharacterized membrane protein RarD (COG2962). Interestingly, COG1054 and COG1179 represent two tRNA modifying enzymes, which we speculate may improve translation capacity in copiotrophs in the context of competitive growth [54, 55].

**Figure 5.**
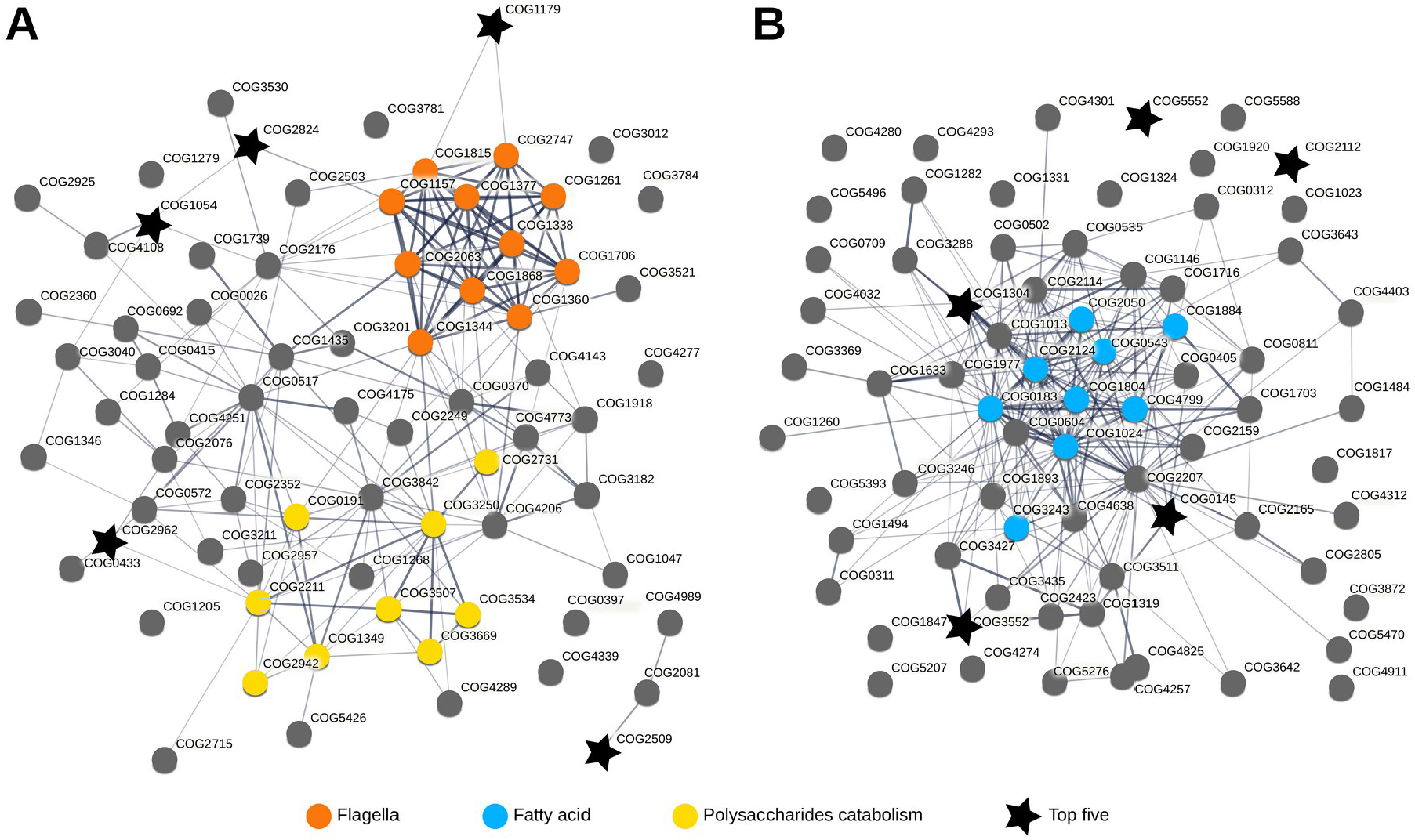
A STRING search of the COGs that were most important for predicting rhizosphere-or soil-association in the RF models, and were enriched in the rhizosphere (A) or in soil (B) according to the PhyloGLM analysis. COGs significantly associated to rhizosphere or to soil were selected from PhyloGLM models, then sorted decreasingly by feature importance in the RF models, and Top 100 most important COGs were selected. Finally, COGs that were associated to both rhizosphere and soil in different taxa in the PhyloGLM models were removed, resulting in 79 COGs uniquely associated with the rhizosphere and 79 uniquely associated with soil. Edge weights represent the level of evidence for functional interaction according to STRING. Some relevant functions are colored according to legend. Top five most important COG features in the RF associated with rhizosphere, or soil according to PhyloGLM are shown.

A similar analysis for soil-associated COGs (Figure 5B) revealed a central cluster that included functions related to fatty acid metabolism, including an Enoyl-CoA hydratase/carnitine racemase (COG1024), a carnitine CoA-transferase (COG1804), an acetyl-CoA acetyltransferase (COG0183), a methylmalonyl-CoA mutase (COG1884), an acetyl-CoA carboxylase (COG4799), a flavin reductase (COG0543), an Acyl-CoA thioesterase (COG2050), cytochrome P450 (COG2124), and a poly-beta-hydroxybutyrate synthase (COG3243). Here, the five most important features in the RF model included a FMN-dependent dehydrogenase (COG1304), an uncharacterized conserved protein (COG3552), a predicted Ser/Thr protein kinase (COG2112), a N-methylhydantoinase A (COG0145), and an uncharacterized protein (COG5552), which are not strongly associated to this central cluster.

The KEGG orthologs (KOs) comprise an alternative functional ontology that we also analyzed. KO-based RF models (Supplementary Table 7) revealed that many of the most important features in rhizosphere bacteria are associated to the “Transporters’’, “Bacterial motility proteins’’, and “Flagellar assembly” pathways, while “Benzoate degradation”, “Aminobenzoate degradation”, and “Glyoxylate and dicarboxylate metabolism” are associated to soil bacteria. One of the top five features associated with soil bacteria is aerobic carbon-monoxide dehydrogenase (K03518), an enzyme involved in metabolizing atmospheric carbon monoxide molecules in biomes with carbon limitation, such as soil [56]. Benzoate degradation is frequently reported in soil bacteria [57], where these abundant aromatic compounds probably are a major carbon source. The glyoxylate cycle may allow soil bacteria to cope with sugar scarcity, as has been shown in another carbon-depleted environment, the marine system [58]. Overall, we hypothesize that rhizosphere-associated bacteria might profit from nutrients coming from root exudates, using expensive flagellar motility and a huge diversity of transporters and enzymes to reach, internalize, and catabolize these compounds, allowing them to reach faster growth rates. In contrast, soil bacteria might use a more costly and slow metabolism involving benzoate, fatty acid degradation, and the glyoxylate pathway to biosynthesize sugars and other carbon compounds in the context of nutrient limitation in the soil.

When we compared our results with previous work by Levy et al. [14] analyzing bacterial traits for root colonization, we found more functions significantly associated either with rhizosphere or with soil bacteria in our analysis, probably because we included a larger set of genomes including both culturable and unculturable bacteria, but also spanning a broader taxonomic range. Approximately 66% of Levy’s significant COGs overlapped with COGs associated with rhizosphere and soil in our analysis (Supplementary Figure 16). Our STRING analysis further revealed that the COGs identified herein were functionally similar, suggesting that our analysis both complements and expands on the previous study by identifying hundreds of new COGs involved in rhizosphere competence.

## Discussion

Understanding rhizosphere microbiomes is critical for microbiome-based crop improvement strategies aimed at crop productivity, plant stress resistance, and soil health. In this work we analyzed rhizosphere and soil data from 16S rDNA metabarcoding studies, and from isolated complete genomes and metagenome-assembled genomes (MAGs), and demonstrated that bacteria enriched in the rhizosphere have higher growth rate potential (lower predicted minimum doubling times, PMDT) than those in soils. This observation holds true in eleven of the most abundant phyla - with the only exception of Firmicutes - independently of host plant genotype, stress condition, soil type, light cycle, or life stage of the host plant. Thus, growth rate potential is a general and important determinant of bacterial rhizosphere competence. Using machine learning models that classify MAGs as being associated with rhizospheres or soils, we could obtain models with high classification accuracy, and revealed that the most important feature for classification is the minimal doubling time. Other important features included several different functions known to be associated with rhizosphere colonization or copiotrophic lifestyle such as flagella, sugar and polysaccharide degradation, and transporters, but also a range of novel protein functions that may be further explored in studies of rhizosphere competence of specific bacteria. Most of the important features associated with soil bacteria included functions related to degradation of aromatic compounds, and the glyoxylate cycle, which may be important to overcome the nutrient limitation in bulk soil. The fact that growth rate potential is the most important determinant explaining rhizosphere microbiome presence is consistent with the notion that the nutritional gradients generated by plant root exudates provide a selective environment for a subset of copiotrophic bacteria from the vast microbial diversity present in soils. Importantly, PMDT can be predicted from the genome sequence, and may thus form a readily accessible feature to predict the rhizosphere competence of bacterial groups that is valid across different plants. Moving forward, determining how much of the growth rate potential is actually realized in specific cases will require careful experimental tracking of individual players in specific plant-microbe systems. As we have shown here, bacterial growth rate potential is one of the most important physiological factors determining their rhizosphere competence and a general feature of rhizosphere-enriched bacteria.

## Acknowledgments

This study was supported by NWO Groen II project number ALWGR.2017.002, the European Research Council (ERC) Consolidator grant 865694: DiversiPHI, and the Deutsche Forschungsgemeinschaft (DFG, German Research Foundation) under Germany’s Excellence Strategy – EXC 2051 – Project-ID 390713860.

