## Supplementary Figures for "Copiotrophs dominate rhizosphere microbiomes and growth rate potential is a major factor explaining the rhizosphere effect"

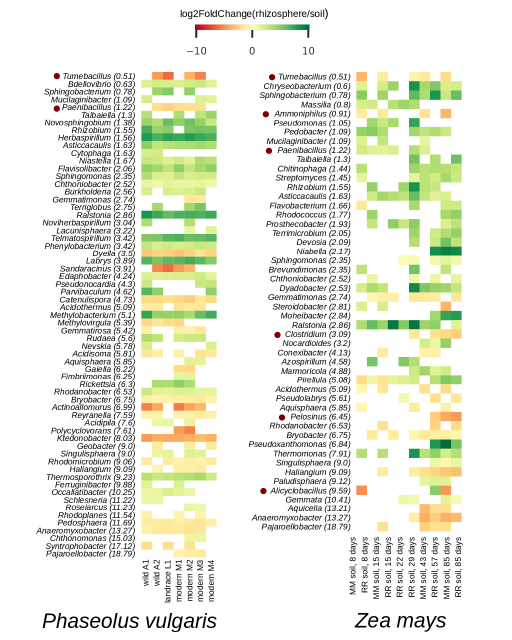

**Supplementary Figure 1. Rhizosphere enrichment in wild and modern accessions of *Phaseolus vulgaris* (common bean), and in *Zea mays* grown in soils with different crop rotation systems.** Differentially abundant genera between rhizosphere and bulk soil replicates were identified by DESeq2 analysis (p-adj < 0.05), and shown here as being significant at least in one experiment. Genera are sorted from lower PMDT (copiotrophs, top) to higher PMDT (oligotrophs, bottom). Numbers in brackets indicate PMDT obtained from the EGGO database for each genus. Brown dots represent genera belonging to the Firmicutes phylum.

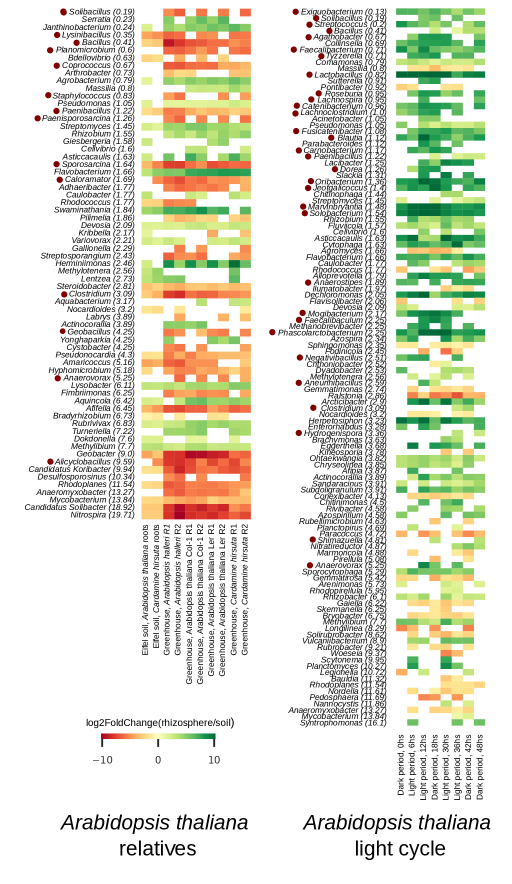

**Supplementary Figure 2. Rhizosphere enrichment in *Arabidopsis thaliana* ecotypes and sister species, and *Arabidopsis thaliana* Col-0 ecotype under light-dark cycle.** Differentially abundant genera between rhizosphere and bulk soil replicates were identified by DESeq2 analysis (p-adj < 0.05), and shown here as being significant at least in one experiment. Genera are sorted from lower PMDT (copiotrophs, top) to higher PMDT (oligotrophs, bottom). Numbers in brackets indicate PMDT obtained from the EGGO database for each genus. Brown dots represent genera belonging to the Firmicutes phylum.

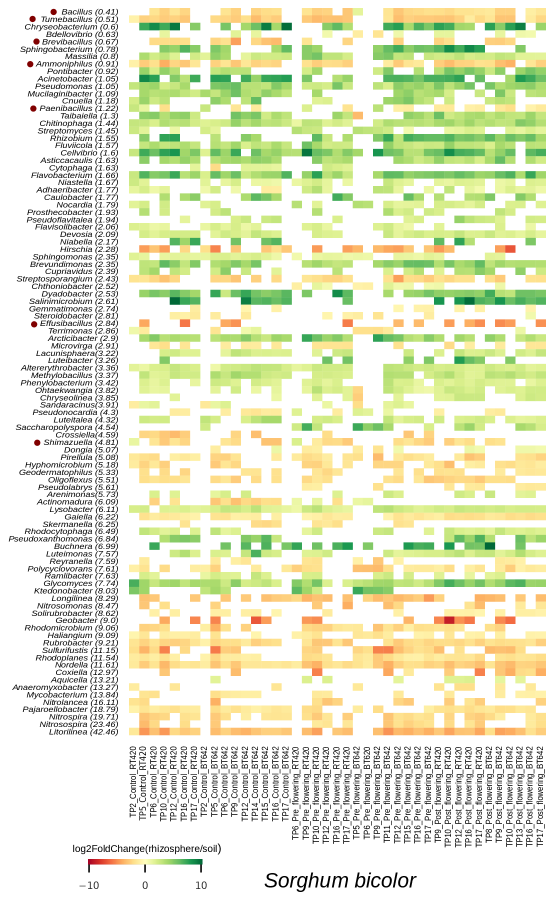

**Supplementary Figure 3. Rhizosphere enrichment in *Sorghum bicolor* under drought stress and control conditions, at different timepoints in its lifecycle.** Differentially abundant genera between rhizosphere and bulk soil replicates were identified by DESeq2 analysis (p-adj < 0.05), and shown here as being significant at least in one experiment. Genera are sorted from lower PMDT (copiotrophs, top) to higher PMDT (oligotrophs, bottom). Numbers in brackets indicate PMDT obtained from the EGGO database for each genus. Brown dots represent genera belonging to the Firmicutes phylum.

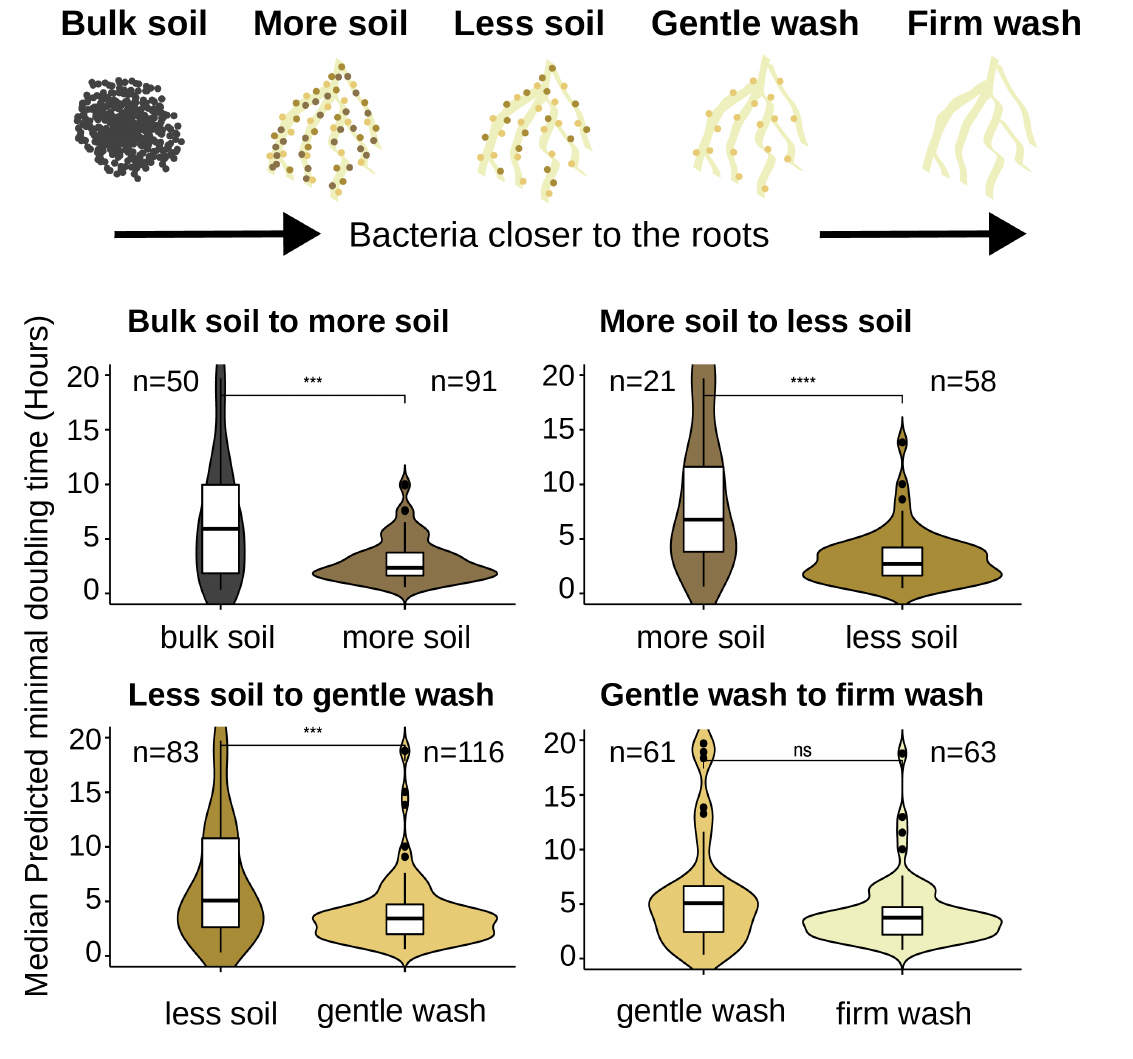

**Supplementary Figure 4. PMDT distributions of genera enriched in *Arabidopsis thaliana* Col-0 sampled at different stages along a bulk soil-to-rhizosphere gradient.** Differentially abundant genera between each pair of the different compartments in the bulk soil-to-rhizosphere gradient were identified by DESeq2 analysis (p-adj < 0.05). PMDT of genera enriched in each compartment are represented. Stepwise compartments from bulk soil to rhizosphere were labeled based on the experimental design in Poppeliers et. al [45]. BS: bulk soil, MS: more soil, LS: less soil, GW: gentle wash, FW: firm wash.

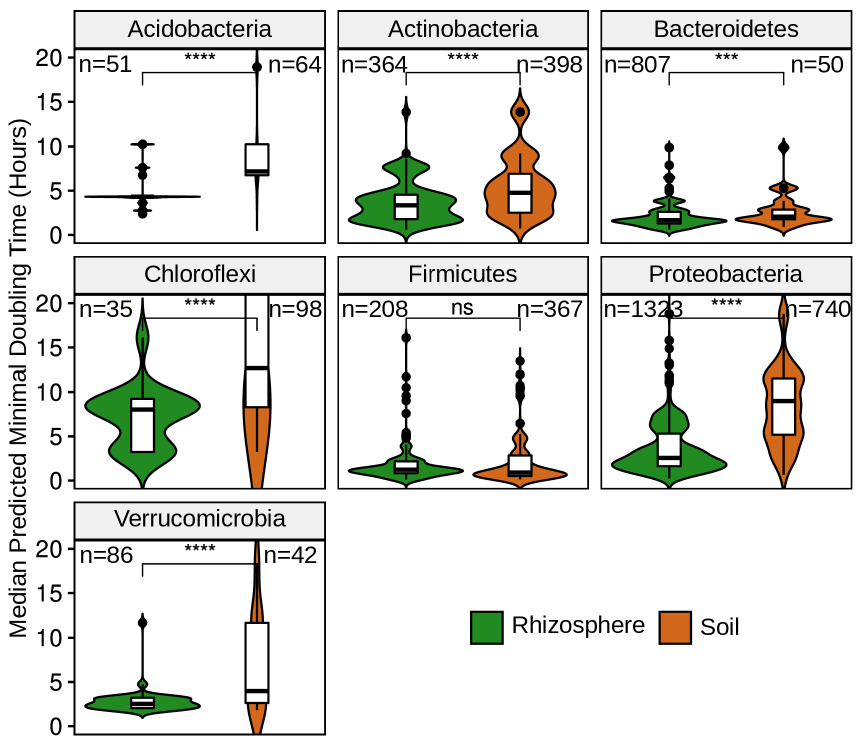

**Supplementary Figure 5. PMDT in rhizosphere and soil enriched-genera by phylum.** Significant enriched genera in all projects were merged and classified in phylum. Violin plots represent those belonging to the most representative phyla. Rhizosphere enriched genera present lower PMDT in Acidobacteria, Actinobacteria, Bacteroidetes, Chloroflexi, Proteobacteria and Verrucomicrobia (Wilcoxon test, p-values < 0.05), but not in Firmicutes.

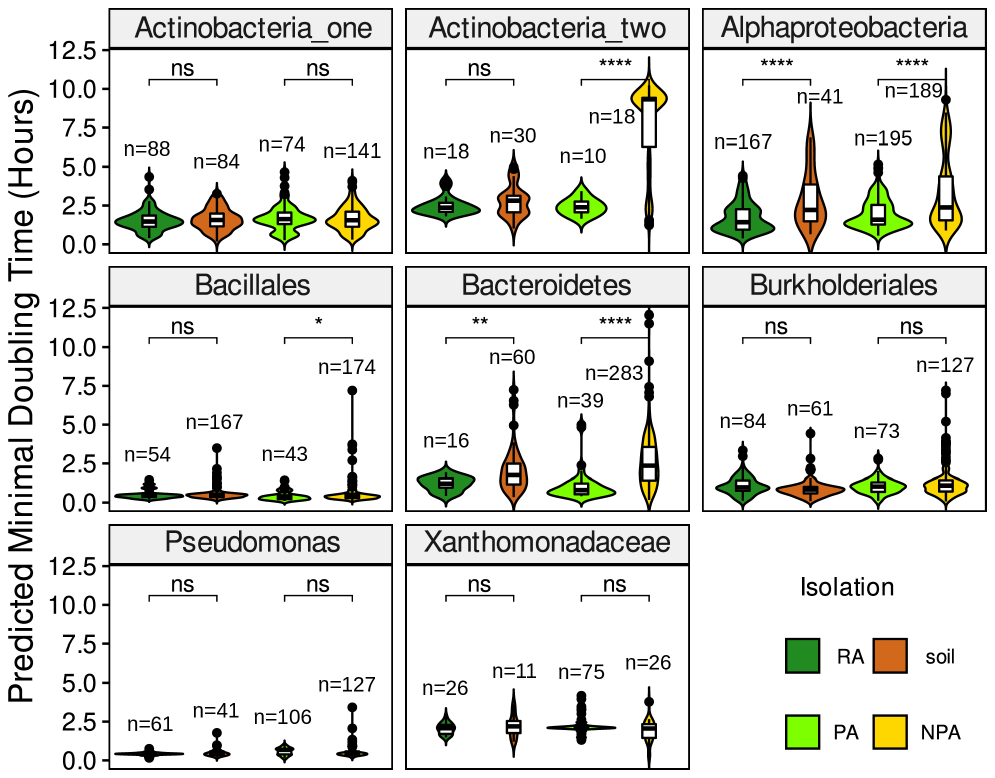

**Supplementary Figure 6: PMDT in Levy et. al. culturable genomes.** Significant differences between genomes isolated from rhizospheres and soils can only be observed in Alphaproteobacteria and Bacteroidetes groups. Distributions of predicted minimal doubling times in genomes from rhizosphere and soils were compared with Mann-Whitney test (ns: p > 0.05, *: p <= 0.05, **: p <= 0.01, ***: p <= 0.001, ****: p <= 0.0001). **RA**: root-associated environments (rhizoplane and endosphere); **PA**: plant-associated environments (including root-associated, and rhizosphere bacteria); **NPA**: non-plant-associated environments, including humans, non-human animals, air, sediments, and aquatic environments; **soil**: bacteria isolated from soils.

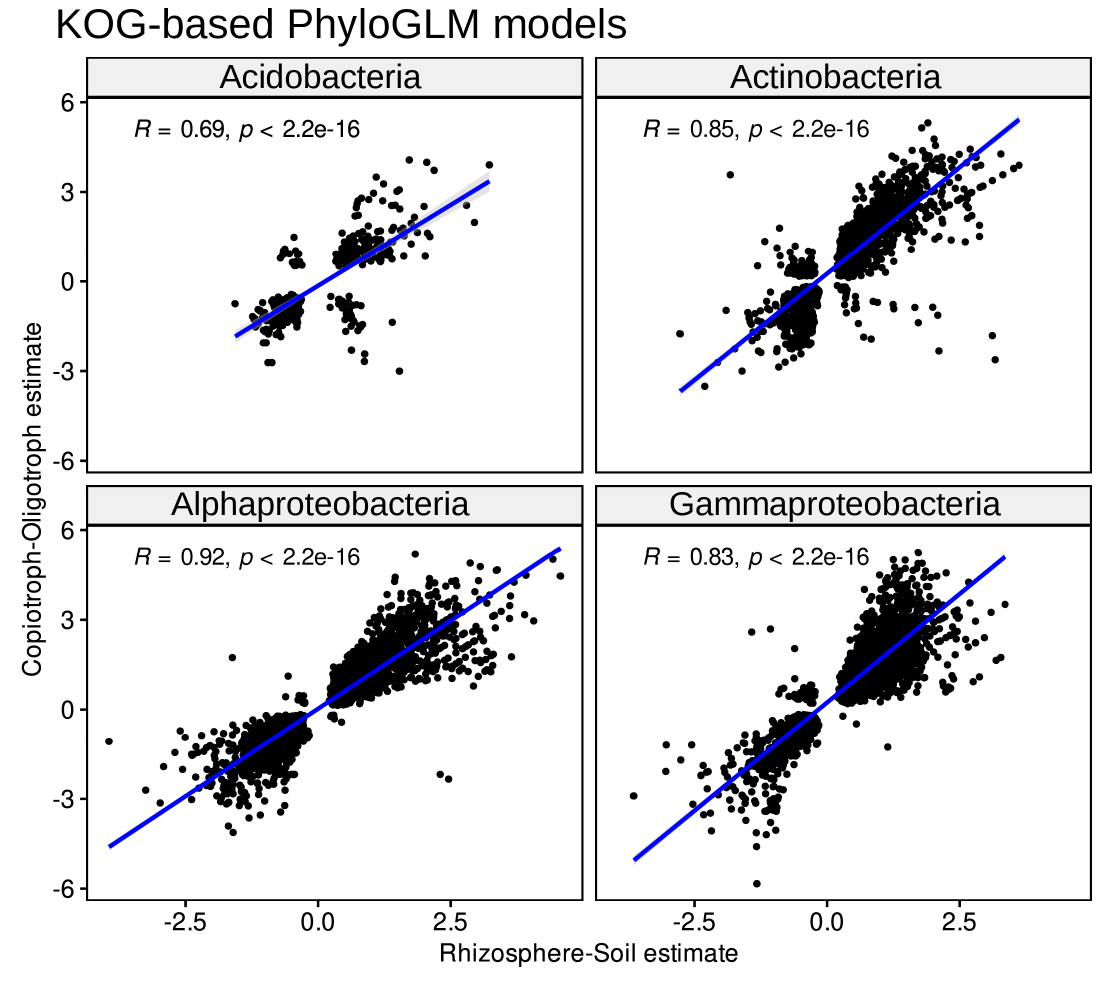

**Supplementary Figure 7. KO-based PhyloGLM estimates correlation between rhizosphere-soil and copiotroph-oligotroph models.** Significantly enriched KO functions in both models (Rhizosphere-Soil and Copiotroph-Oligotroph, PhyloGLM FDR < 0.05) positively correlate in Alphaproteobacteria, Gammaproteobacteria, Acidobacteria, and Actinobacteria, indicating that most functions enriched in rhizosphere MAGs are also enriched in copiotroph MAGs, and vice versa. Spearman correlation tests are shown.

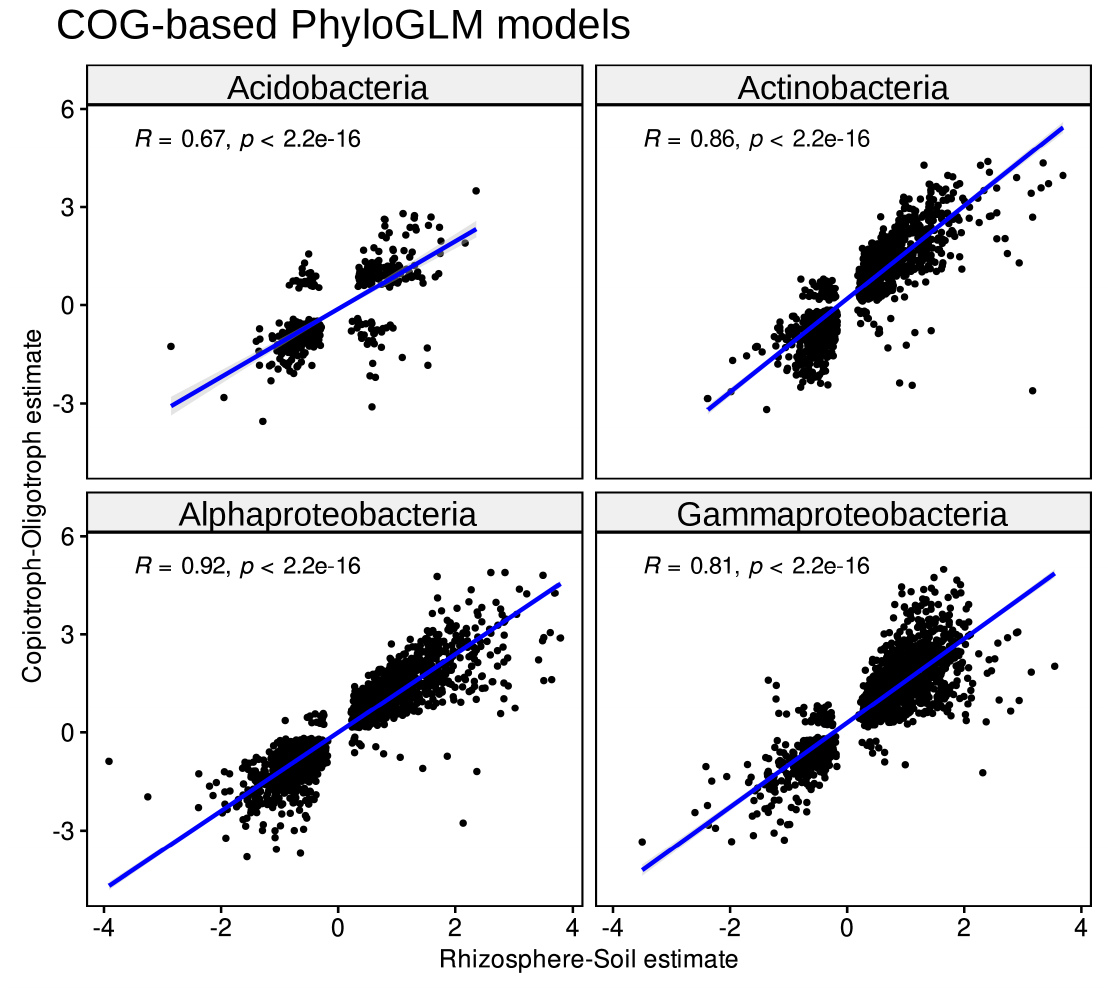

**Supplementary Figure 8. COG-based PhyloGLM estimates correlation between rhizosphere-soil and copiotroph-oligotroph models.** Significantly enriched COG functions in both models (Rhizo-Soil and Cop-Oli, PhyloGLM, FDR < 0.05) positively correlate in Alphaproteobacteria, Gammaproteobacteria, Acidobacteria and Actinobacteria, indicating that most functions enriched in rhizosphere MAGs are also enriched in copiotroph MAGs, and vice versa. Bacteroidetes show no correlation. Spearman correlation tests are shown.

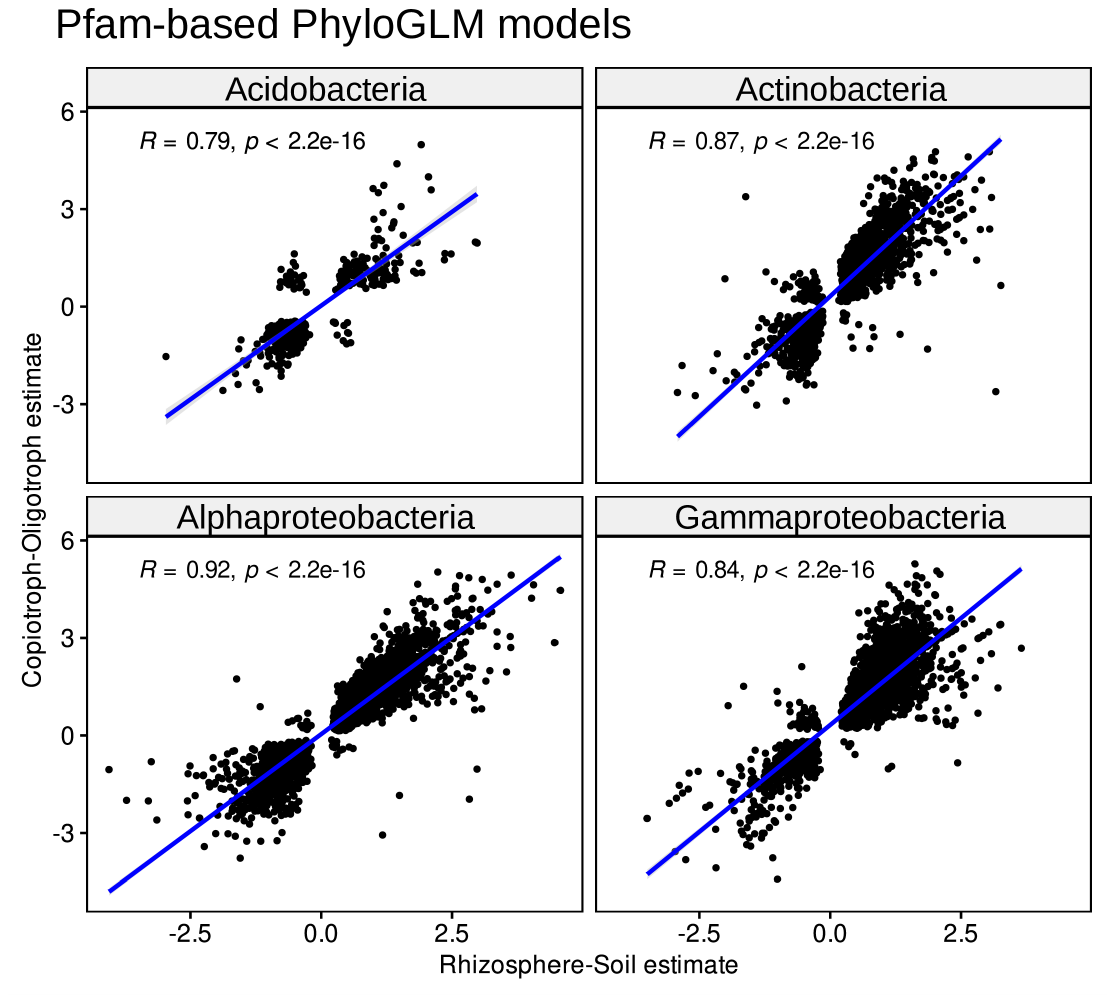

**Supplementary Figure 9. Pfam-based PhyloGLM estimates correlation between rhizosphere-soil and copiotroph-oligotroph models.** Significantly enriched Pfam functions in both models (Rhizo-Soil and Cop-Oli, PhyloGLM, FDR < 0.05) positively correlate in Alphaproteobacteria, Gammaproteobacteria, Acidobacteria and Actinobacteria, indicating that most functions enriched in rhizosphere MAGs are also enriched in copiotroph MAGs, and vice versa. Spearman correlation tests are shown.

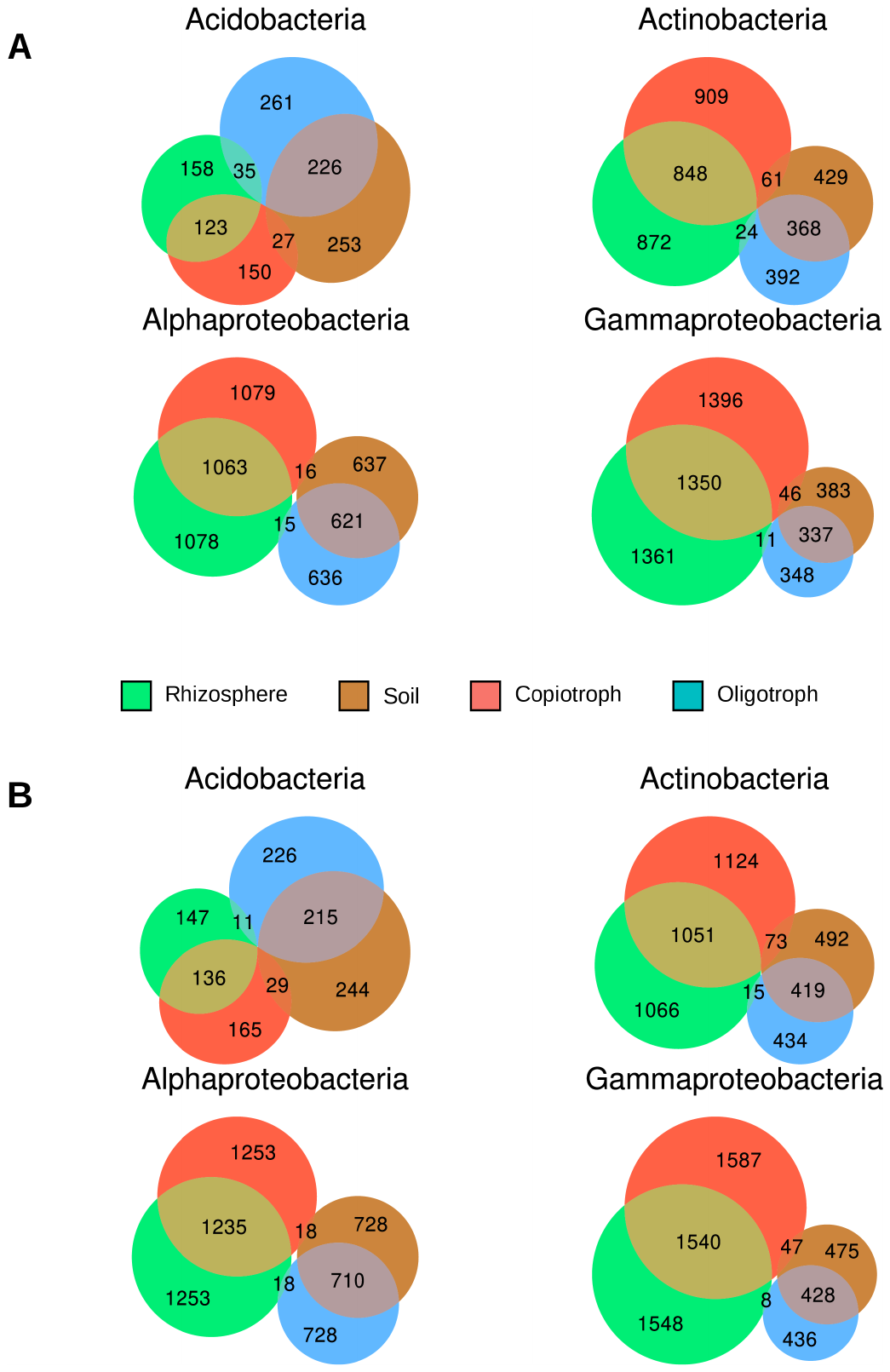

**Supplementary Figure 10: Euler plots with significant functions enriched in MAGs.** Number of enriched functions in PhyloGLM models in rhizospheres (green) or soil (brown), and in copiotrophs (red) or oligotrophs (blue). Functions mainly enriched in MAGs from rhizospheres are also the same being enriched in copiotrophs, similarly functions enriched in soils are mainly enriched in oligotrophs. A. COG-based PhyloGLM models. B. Pfam-based PhyloGLM models.

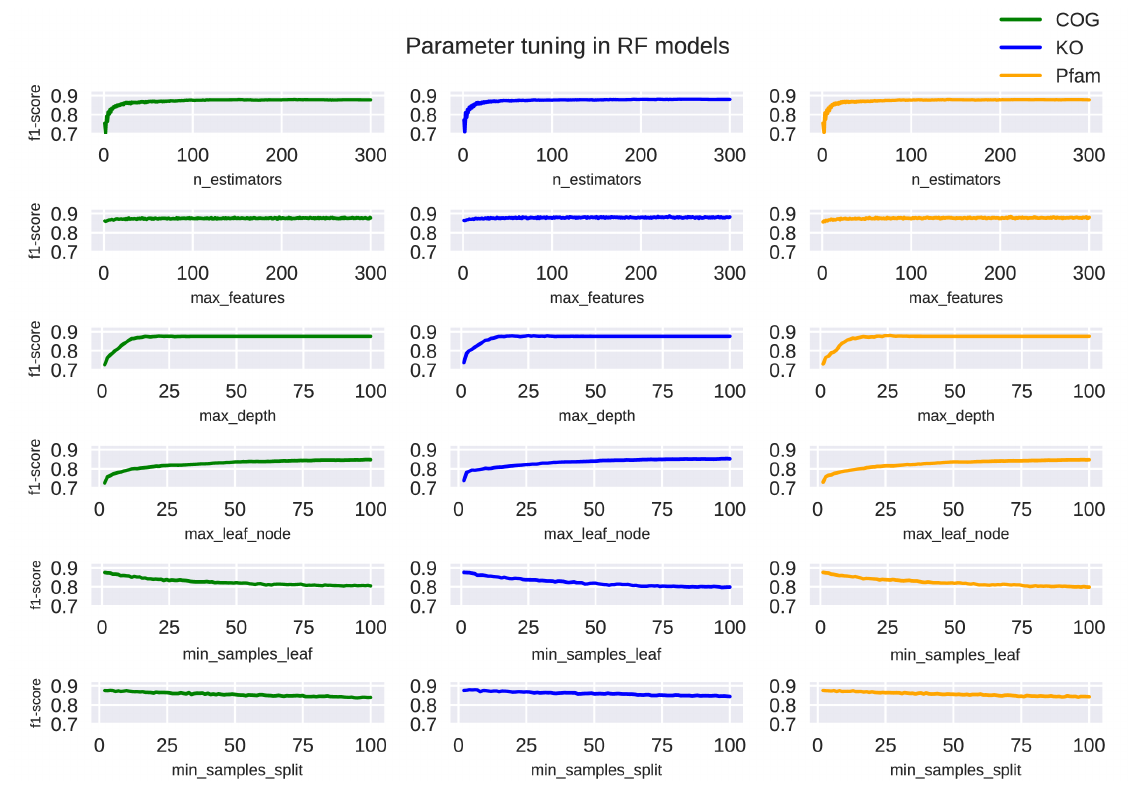

**Supplementary Figure 11: Parameter tuning for Random Forest models.** RF classification models were trained using COG, KO and Pfam binary matrices with copiotroph / oligotroph as an additional dummy feature.

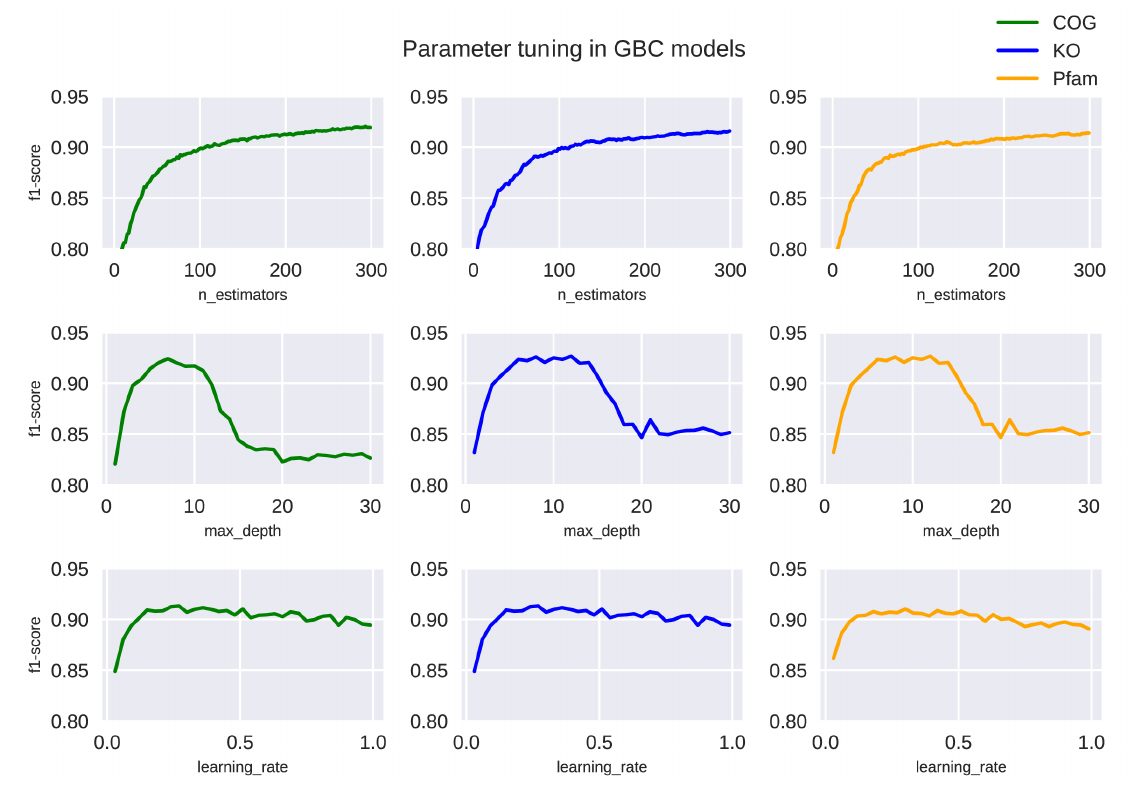

**Supplementary Figure 12: Parameter tuning for Gradient Boosting Classifier models.** Parameters n_estimators, max_depth, and learning_rate were changed, and f1-scores were used as criteria to define optimal ranges for these parameters.

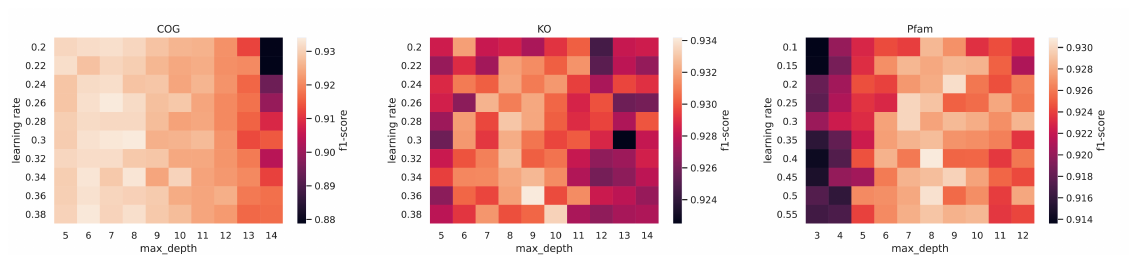

**Supplementary Figure 13**: F1-scores of a grid search combining max_depth and learning_rate parameters in GBC COG (A), KO (B) and Pfam (C) models, with n_estimators=300. Optimal parameters were max_depth=8, 9, and 8, and learning_rate=0.30, 0.36, and 0.40, respectively.

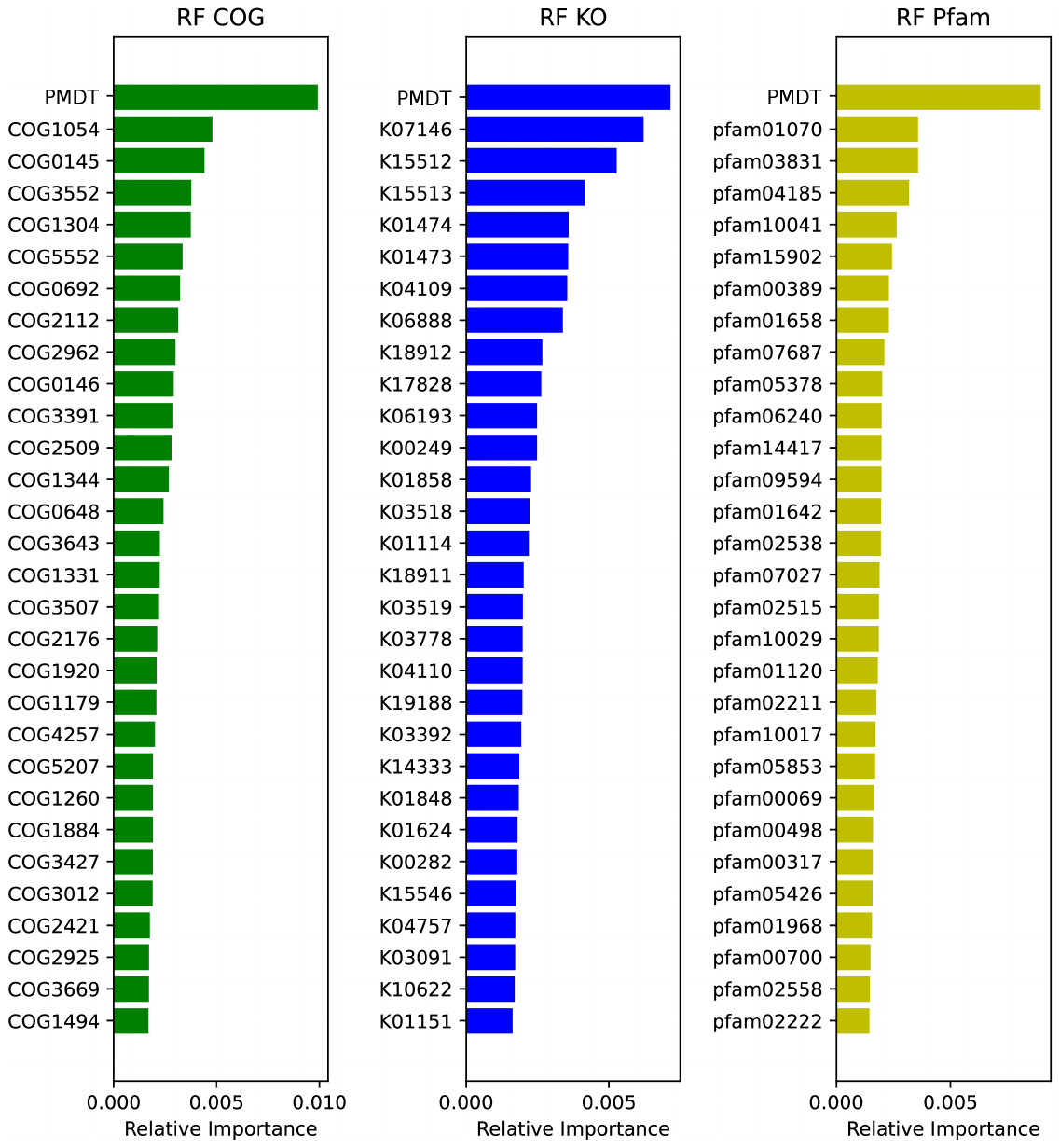

**Supplementary Figure 14. Feature importances of RF models based on COG, KO, and Pfam.** Models were trained with optimal parameters and their 30 most important features are displayed.

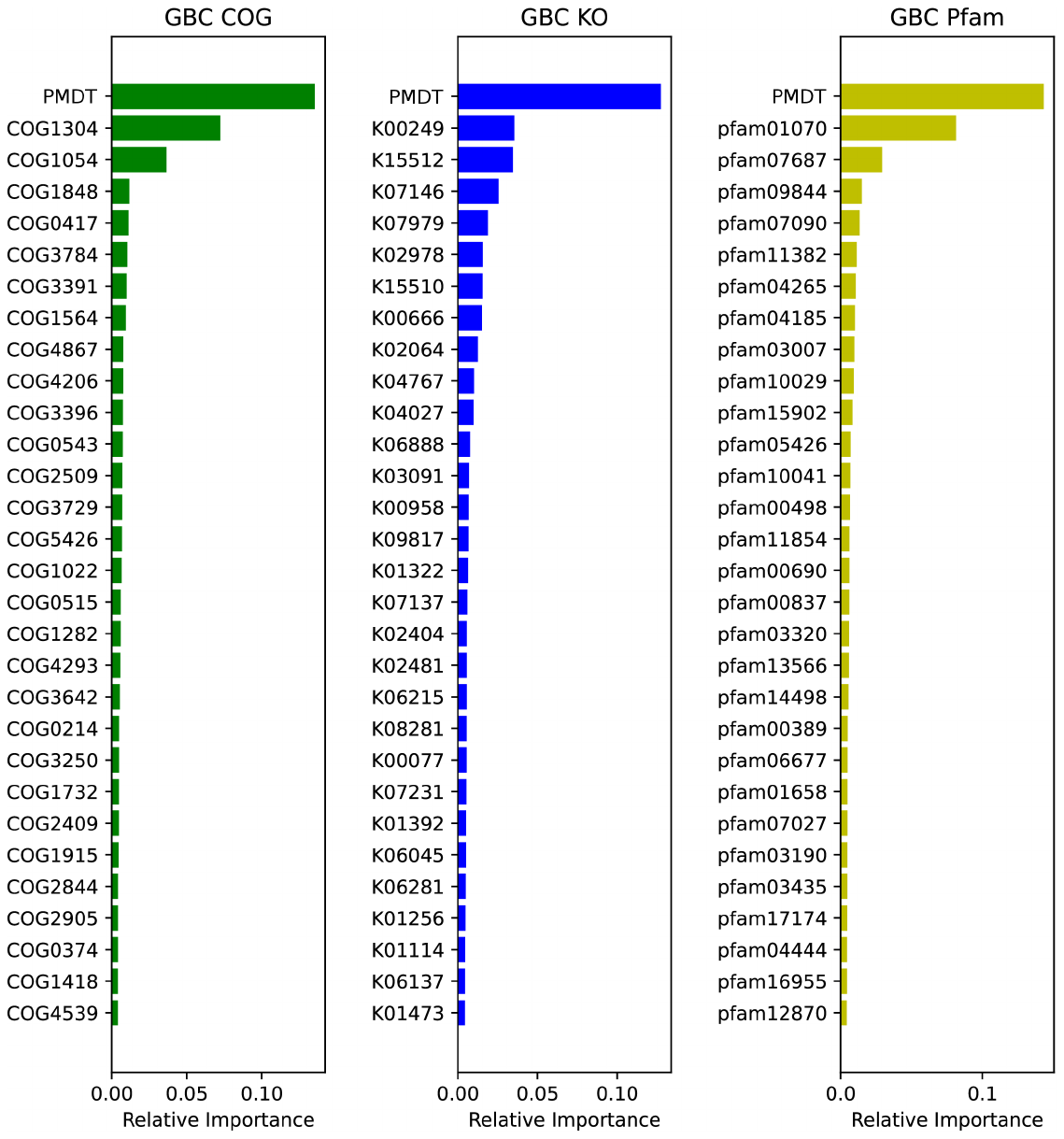

**Supplementary Figure 15. Feature importances of GBC models based on COG, KO, and Pfam.** Models were trained with optimal parameters and their 30 most important features are displayed.

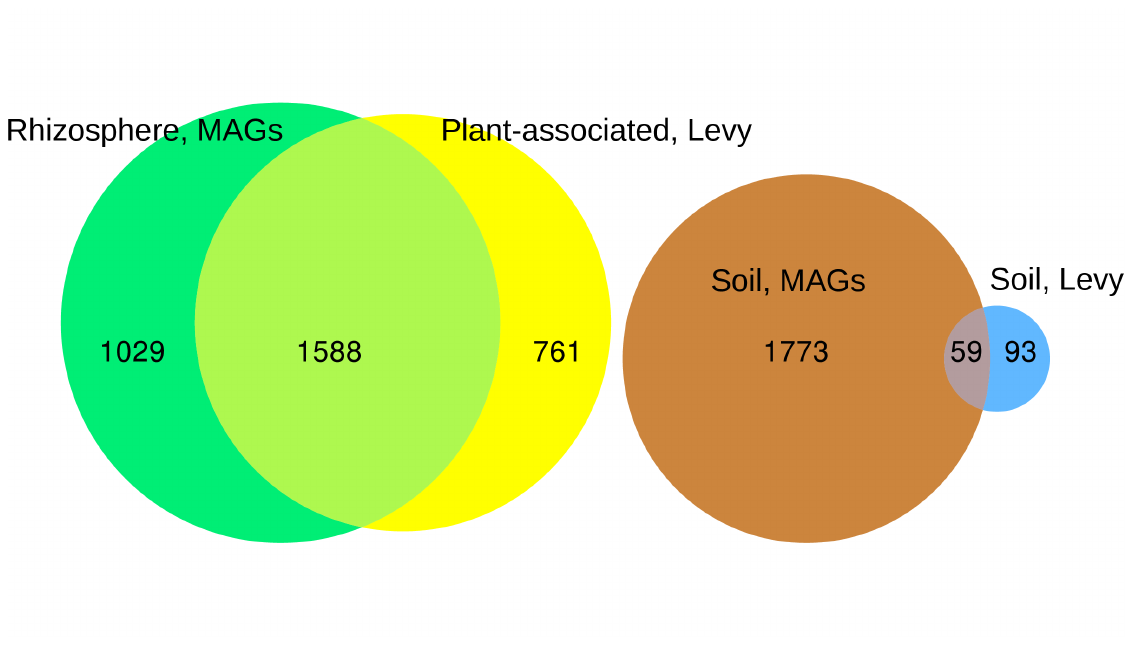

**Supplementary Figure 16. Significant COGs found in our work and in Levy et al.** Numbers correspond to COGs found significantly associated with rhizosphere and soil bacteria in our work (green and brown) and in Levy's work (yellow and blue), respectively. From 41240 COG-based PhyloGLM tests performed in our work (all COGs in each of the 11 taxonomic groups defined here), 2617 COGs were uniquely associated to rhizosphere MAGs, and 1832 uniquely to soil MAGs. Similarly, numbers from Levy's work correspond to COGs significantly associated to uniquely plant-associated genomes or uniquely to soil genomes with any of the 5 statistical tests done by the authors, with 2349 COGs associated to plant-associated genomes, and 152 associated to soil genomes.
